# A switch in the mode of tissue growth extends the growth phase of *Drosophila* wing primordia during early pupal development

**DOI:** 10.1101/2025.01.09.632245

**Authors:** Khaoula El Marzkioui, Isabelle Gaugué, Ettore De Giorgio, Pierre Léopold, Laura Boulan

## Abstract

Understanding how final organ size is established during development raises two key questions: how organs increase their mass and how they stop growing upon reaching an appropriate size. While organ growth is driven by conserved signaling pathways, the mechanisms underlying growth arrest remain elusive. Studies on *Drosophila* imaginal wing discs have provided a model whereby final organ size coincides with an arrest of cell proliferation at the end of the larval phase. Here, through 3D reconstruction and volume measurements, we show that wing discs grow continuously throughout the larva-to-pupa (L/P) transition and proceed to growth arrest later during the pupal period. This supplemental growth phase involves a switch at the L/P transition that uncouples proliferation from tissue growth, with an important contribution from Insulin/IGF signaling. These findings challenge the existing model of imaginal wing development and open new avenues for our understanding of growth arrest and organ size determination.

## INTRODUCTION

The determination of organ size is a complex process that relies on the integration of autonomous and non-autonomous signals. Autonomous growth programs, primarily driven by morphogens, play a crucial and conserved role in tissue patterning and growth regulation^1^. Meanwhile, systemic signals adjust organ growth in response to environmental factors, such as nutrition, as well as developmental cues^2^. This coordination ensures species-specific organ shapes and proportions. The study of growth control encompasses two interrelated questions: what drives growth, and what stops it. Although conserved signaling pathways that regulate tissue growth are now well-characterized, the cellular and molecular mechanisms that underlie growth arrest are far less understood.

Over the past few decades, organ transplantation and regeneration experiments provided general concepts regarding organ size determination^3,4^. Research on *Drosophila* imaginal discs has started entangling the complex integration of growth control with developmental processes^5–7^. During larval stages, the wing disc is a highly growing and proliferating monolayer epithelium. Towards the end of larval development, cell proliferation slows down and finally stops at the larva-to-pupa (L/P) transition. In early pupa, wing cells are arrested at the G2 stage^8,9^. A final wave of 2-3 cleavage (size-reductive) divisions starts 12-16 hours after pupa formation (APF), culminating in a final G1 cell cycle exit around 24h APF^9–11^. Growth has traditionally been associated with proliferation, leading to a model whereby proliferation arrest determines final tissue size at the L/P transition. This is supported by measurements of tissue area suggesting a sharp decrease in disc expansion rate at the end of the third larval instar^12–14^. Consequently, the L/P transition is believed to mark a temporal switch between growth and morphogenesis.

Understanding proliferation arrest at the L/P transition has been the focus of both experimental and theoretical studies. “Mechanical feedback” models for uniform proliferation and growth arrest propose that local morphogen-driven growth promotes tissue compression, which in turn exerts negative feedback on cell proliferation^15–17^. At the systemic level, high titers of the steroid hormone ecdysone at the L/P transition contribute to G2 arrest in wing disc cells via inhibition of *string/cdc25* expression^10^. Additionally, TOR signaling integrates various inputs during late larval development, including the effects of ecdysone on cell proliferation^18^ and the growth-inhibitory effect of developmental hypoxia^19^.

In this study, we revisit wing disc growth by measuring tissue volume as a proxy for tissue mass. Contrarily to previous experimental descriptions and models, we observed that the volume of wing primordia increases continuously during larval and early pupal stages. Consistent with proliferation arrest at the L/P transition, pupal wing growth is driven by cell volume increase only. These findings suggest that final wing tissue mass is determined during the pupal phase, independently of cell proliferation arrest, revealing a new phase of tissue growth that takes place during wing eversion, expansion and elongation. We analyzed the role of classical growth-regulatory pathways and highlighted a crucial contribution of Insulin/IGF signaling in pupal wing growth, despite the animals being in a nonfeeding stage. In conclusion, our findings redefine the time window for growth arrest and highlight the existence of an additional growth phase in the early pupal stage.

## RESULTS

### Growth arrest occurs during the pupal stage

Our first objective was to define more precisely when growth arrest occurs in the wing primordium. The current model of growth arrest at the L/P transition comes from the use of cell proliferation as a marker of tissue growth, but also from the difficulty to quantify tissue size during the early pupal stage. Indeed, the L/P transition marks the beginning of important morphogenetic events starting with tissue eversion, which strongly perturbs anatomical landmarks. It is therefore not possible to compare tissue area in wing primordia as 2D structures (obtained from projections or cross-sections along the proximo-distal axis) before and after tissue eversion. To bypass this limitation, we decided to use 3D reconstruction of wing primordia to quantify tissue volume. This experimental approach has previously allowed us to quantify fluctuating asymmetry with high precision during development^20^. To establish the growth trajectory of the wing primordia, we expressed a *UAS-GFP* transgene under the control of the wing-specific *rotund-GAL4* driver (*rn-GAL4*), which allowed us to perform 3D reconstruction both during larval and pupal development (Figure 1A). For this purpose, we synchronized animals twice: first at L1 larva hatching, and later at the L/P transition (which occurs at 124h AED in *rn>GFP* animals, Figure S1A). Using this setup, we measured tissue volume and observed that wing growth persists after the L/P transition and until 16h after pupa formation (APF), after which the volume of the pupal wing remains constant. Noticeably, wings increase their volume around 2.5-fold after the L/P transition (Figure 1B). Calculation of the growth rate based on tissue volume reveals a slow decline during the last larval stage, which was recently shown to correlate with physiological hypoxia^19^. However, the growth rate stabilizes for approximately 12-16h after the L/P transition, before its final drop reflecting growth arrest (Figure 1C).

**Figure 1:**
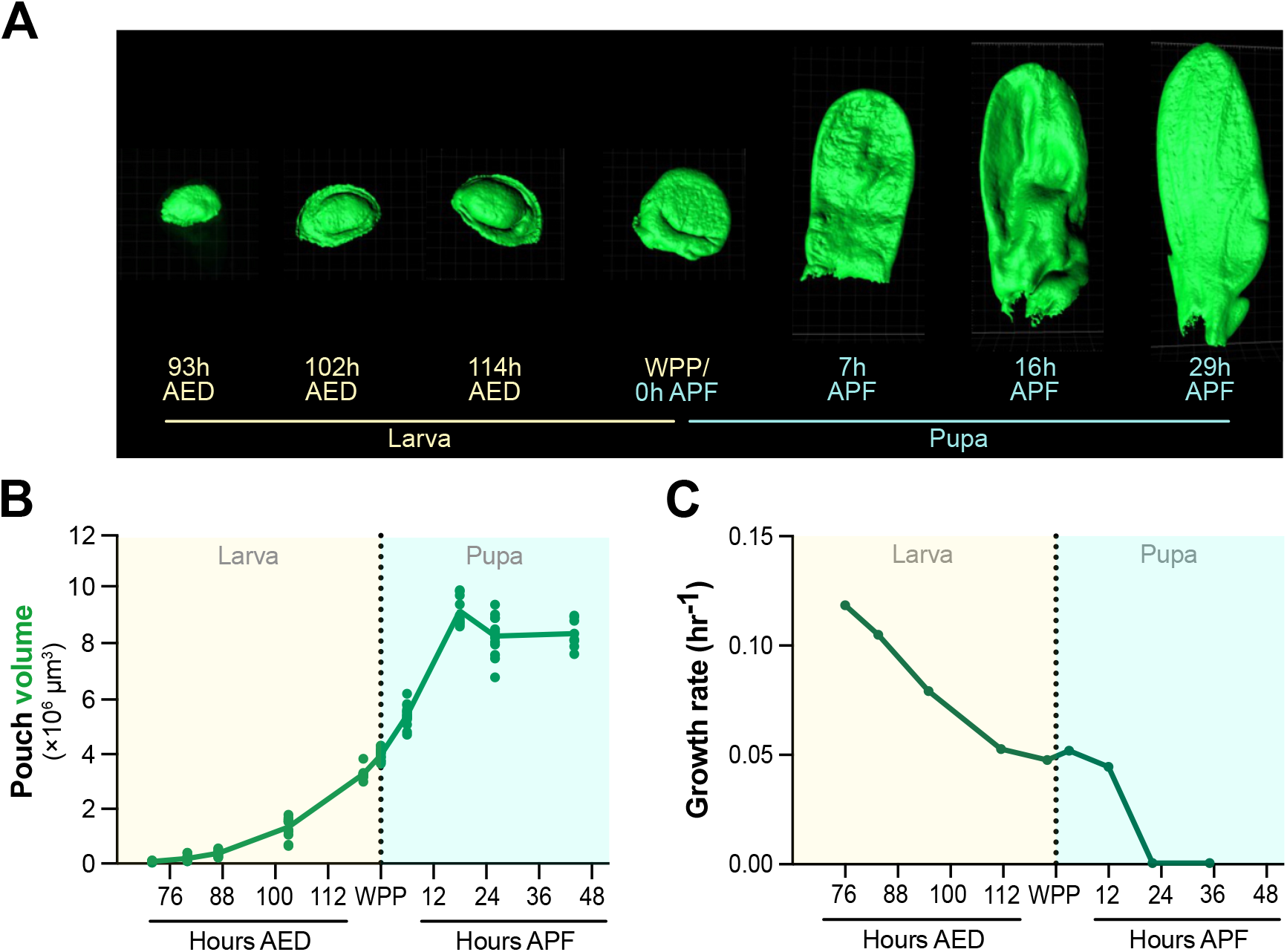
Measuring tissue volume reveals that wing growth extends during the early pupal phase. A) Representative examples of 3D reconstruction of wing primordia expressing *UAS-GFP* under the control of the *rn-GAL4* driver. All stages are shown with the same magnification. B) Growth curve based on volume measurement of the GFP-expressing domain at the indicated timepoints (only females were considered). C) Growth rate calculations based on the data presented in B. AED: After egg deposition; APF: After pupa formation; WPP: White Prepupa, corresponds to 0h APF. The dashed line represents the larva-to-pupa transition.

As a matter of comparison, we used the same samples projected in 2D to measure the area of the GFP-labelled wing pouch and the corresponding growth rate during the third larval instar. Our data fit with previously published datasets on wing disc area during larval development^12–14^. Therefore, tissue area slows down more drastically at the end of larval development than does tissue volume (Figures S1B-C). This discrepancy can be attributed to the thickening of the epithelium during the late 3^rd^ instar^21,22^ and emphasizes the importance of considering volumes as a proxy for mass rather than area, even in the case of a monolayered epithelium.

Importantly, when compared with wing morphological markers like the expression domain of the *wingless* gene, *rn-GAL4* expression domain is precisely conserved between larval and pupal stages (Figure S1D), confirming that it can faithfully be used for delimiting the wing primordium during development. *nab-GAL4*, another wing driver active during the pupal stage, gave similar results and confirmed the increase in tissue volume in early pupae (Figure S1E). Altogether, these results unravel a previously unknown phase of tissue growth and indicate that growth arrest of wing precursors takes place during the pupal stage.

### Cell growth sustains tissue growth in the early pupa

To investigate how wing primordia grow in the early pupa, we decided to quantify cellular parameters. First, we took advantage of staining the cells for fibrillarin, which distinctly labels each cell with a single dot in the nucleolus (Figure S2). This allowed us to estimate the total number of cells within the wing pouch and calculate the average cell volume. Consistent with the proliferative phase ending around the L/P transition, cell number increases during the larval stage until reaching a plateau at the L/P transition (Figure 2A). Conversely, cell volume is rather constant during the larval stage, reflecting the coupling between cell growth and cell divisions, but drastically increases during the early pupal stage (0-16h APF). At 29h APF, cell number raises again while cell volume drops, in agreement with the final size-reductive cell divisions taking place during the 16-24h APF time window^9,11^. We then induced small clones (2-4 cells) marked with membrane GFP to perform measurements at cellular scale during the 0-17h APF period. This approach further supports that early pupal wing growth is driven by an increase in cell volume (Figure 2C).

**Figure 2:**
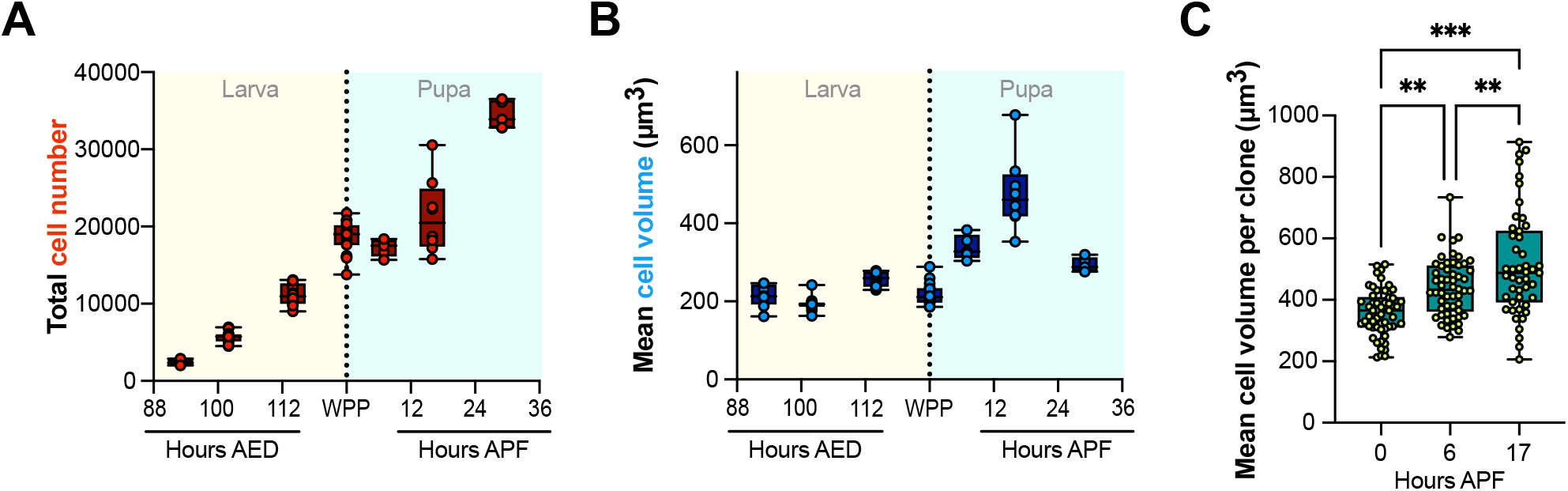
Wing growth during the early pupal stage relies on an increase in cell volume. A) Average cell volume at the indicated developmental times. B) Estimated total cell number. C) Quantification of average cell volume in clones (2-4 cells) marked with PH-GFP at the indicated time points. AED: After egg deposition; APF: After pupa formation; WPP: White Prepupa, corresponds to 0h APF. The dashed line represents the larva-to-pupa transition. Box and whisker plots display min to max values and individual datapoints.

### Modulating growth pathways in the early pupal stage aBects final adult wing area

We next undertook to characterize which growth-regulatory pathways affect wing development during early pupa. Using an inducible expression system (*nub-GAL4, tub-GAL80*^*TS*^ combined with *elav-GAL80* to prevent neuronal expression), we targeted transgenes specifically in the wing pouch during the 0-16h APF time window and analyzed the resulting variations in adult wing area (Figure 3A). Figure 3B displays wing area measurements and Figure S3 provides representative wing images. In this experimental set up, a limiting regulator for growth during the pupal stage would be expected to produce tissue growth reduction when inhibited and, conversely, growth increase when overactivated.

**Figure 3:**
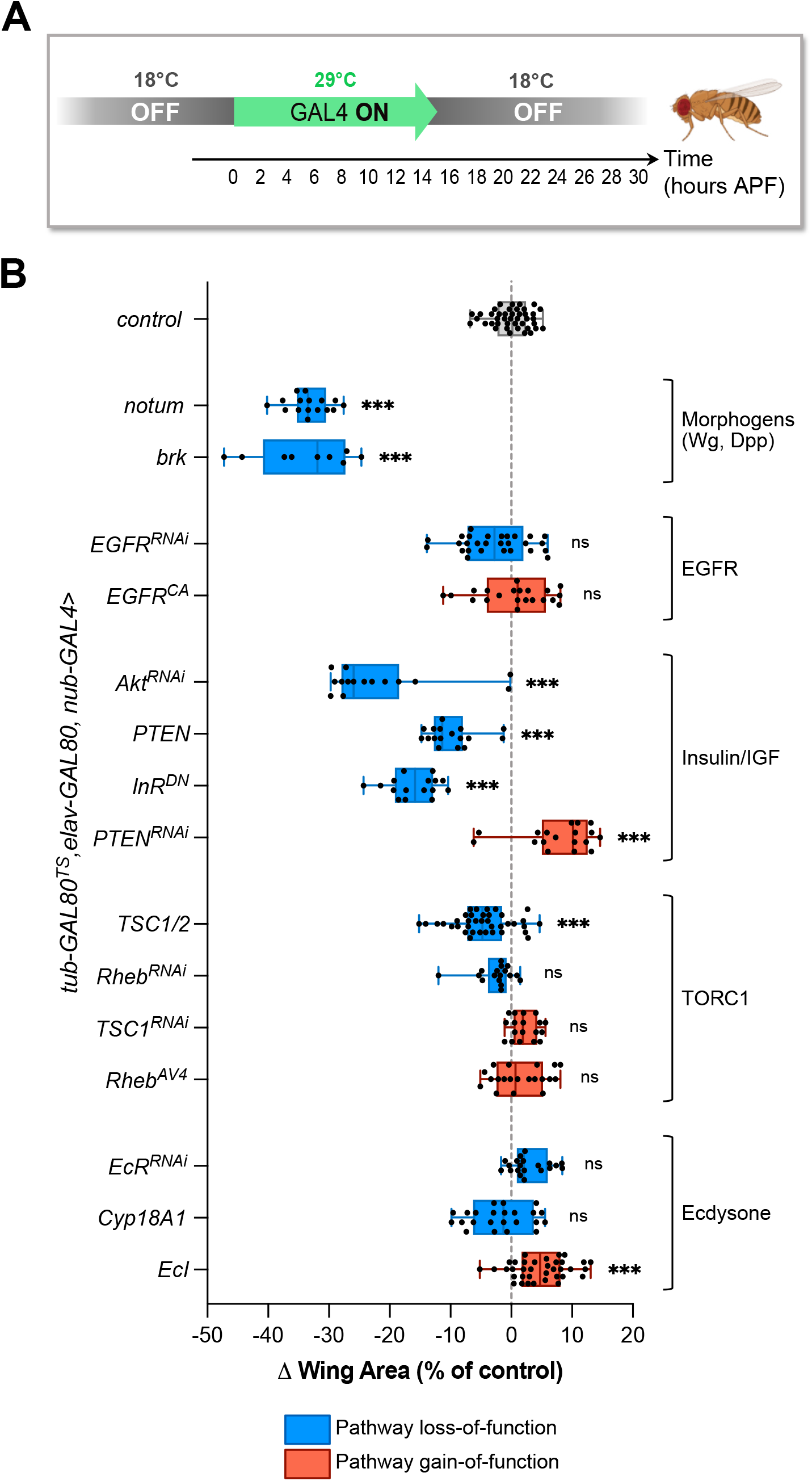
Transient modulation of growth pathways and their eBect on adult tissue size. A) Experimental setup: using the GAL80TS system, GAL4 expression in wing primordia was restricted to a 16h time window at the beginning of the pupal stage. The resulting adult wings were analyzed. B) Ehect of the indicated genetic manipulations on final wing area, shown as the deviation compared to the control (grey dashed line). Box and whisker plots display min to max values and individual datapoints.

Inhibiting the activity of the Dpp and Wg morphogens (by driving expression of their inhibitors: *notum* for Wg signaling and *brinker* for Dpp signaling) in early pupae resulted in a significant reduction in adult wing area and pronounced morphological defects as previously reported^23–25^. Modulations of EGFR activity did not affect wing area and had minimal effects on vein development. We then examined pathways that convey systemic/environmental information. The most dramatic effects were observed when modulating the Insulin/IGF signaling (IIS) pathway. Indeed, blocking insulin receptor activity (*UAS-InR*^*DN*^) or decreasing intracellular signaling (*UAS-Akt*^*RNAi*^ and *UAS-PTEN*) significantly reduced wing area without disrupting tissue patterning. Conversely, increased IIS (*UAS-PTEN*^*RNAi*^) led to significantly larger adult wings. The contribution of the TORC1 pathway appeared milder, as only overexpressing the TSC1/2 inhibitors resulted in a slight reduction in tissue area. Finally, we modulated the response to ecdysone by acting on its receptor (*UAS-EcR*^*RNAi*^) or on intracellular ecdysone concentrations (overexpressing the degradation enzyme Cyp18A1^26^ or the ecdysone importer EcI^27^). While increasing ecdysone import (*UAS-EcI*) in the early pupa resulted in a slight increase in adult wing size, both conditions of decreased ecdysone signaling had no effect on final wing size or morphology. This suggests that, although elevated ecdysone levels can enhance tissue growth, ecdysone signaling is not essential for tissue growth at this stage. Since the metamorphic peak of ecdysone triggers wing eversion at the L/P transition, the absence of obvious morphological phenotypes (Figure S3) indicates that our shift protocol for GAL4 activation has no effect on late larval development.

### Insulin/IGF signaling plays a key role for early pupal wing growth

We have observed that both autonomous (morphogens) and systemic (Insulin/IGF, TOR) signals are important in the early pupa for determining the surface of adult wings. However, adult wings are flat cuticular structures devoid of living cells, whose area is determined by the integration of two main processes: (i) wing tissue growth and (ii) tissue expansion and elongation processes that shape the final cuticular scaffold^28^. To establish whether the phenotypes observed in adult wings reflect the role of signaling molecules in tissue growth proper, we measured tissue volume in mid-pupa after growth arrest. To improve the temporal resolution and the precision of our genetic manipulations, we implemented the recently published ShineGAL4/UAS optogenetic system^29^ for pupal wings by generating a *rotund-ShineGAL4* line (hereafter *rn-ShineGAL4*). This system offers the advantage of inducing gene expression through dark-to-light shifts, thereby avoiding perturbations of developmental timing inherent to the temperature shifts used with the GAL80^ts^ system.

Using *rn-shineGal4*, we modulated the activity of candidate pathways after shifting animals to daylight at the L/P transition (Figure 4A). Pupal wings were dissected and analyzed at 20h APF, shortly after growth arrest. Surprisingly, inhibition of Dpp and Wg signaling had no effect on pupal wing volume (Figure 4B). In contrast, both gain- and loss-of-function for IIS had a significant impact on wing volume at 20h APF, identifying this pathway as a regulator for pupal wing growth. We also confirmed that inhibiting -but not enhancing-TOR signaling affects pupal wing growth, suggesting that the pathway is necessary but not sufficient to sustain growth at this stage. Finally, measurements of tissue volume under ecdysone signaling inhibition corroborated our observations that this hormone is not necessary for wing growth during early pupal stage. Therefore, our results suggest that the role of ecdysone signaling in wing growth is restricted to larval development^30–33^.

**Figure 4:**
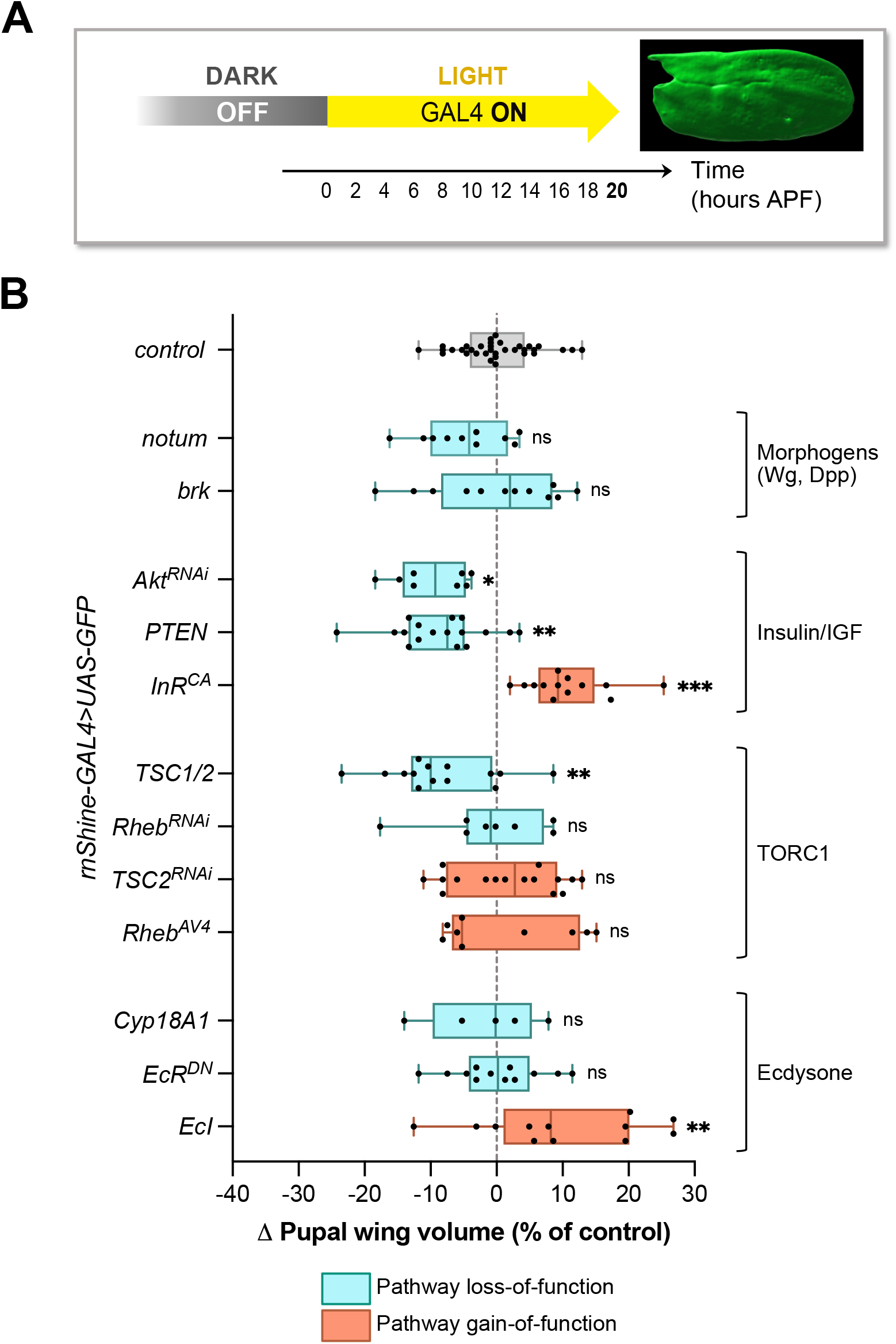
Insulin signaling is a major input for early pupal wing growth. A) Experimental setup: using the Shine-GAL4 system, GAL4 expression was turned on at the L/P transition (dark-to-light shift) to modulate the activity of growth pathways specifically during the early pupal stage until dissection at 20h APF. B) Effect of the indicated genetic manipulations on the volume of pupal wings at 20h APF, shown as the deviation compared to the control (grey dashed line). Box and whisker plots display min to max values and individual datapoints.

## DISCUSSION

Using tissue volume as a proxy for mass, we show here that, contrarily to the current dogma, final wing tissue mass is reached approximately 16 hours after the onset of pupariation. Notably, we observed that the growth rate is constant across the L/P transition, suggesting that proliferation arrest is uncoupled from volume increase at that stage. This revised timing for growth arrest implies that growth and morphogenesis overlap, suggesting that these processes are coupled to ensure proper organogenesis.

Many scenarios of mass increase involve cell proliferation, leading to the conceptual overlap of these two processes, with proliferation often serving as a marker for growth. However, they can be uncoupled in the life history of an organism^34^. Increasing evidence also shows that cell number is not the primary determinant for organ size. Instead, growth control mechanisms seem to integrate higher levels of size sensing. For instance, in diploid and triploid *Xenopus laevis* embryos, cell size scales with ploidy but cell number adjusts to produce individuals with similar body mass^35^. At tissue scale, alterations of the cell division rate in *Drosophila* wing discs are compensated for by changes in cell volume to adjust tissue size^36,37^.

Here we show that organ growth continues after the L/P transition despite a drastic drop in cell proliferation, raising the question of the biological relevance of this cell cycle arrest. One possibility is that proliferation arrest allows cellular physiology (acto-myosin/tubulin networks) to adapt to the drastic morphogenetic processes starting at the L/P transition^28^. The transition from proliferative to non-proliferative growth could thus ensure proper coordination between tissue mass increase and morphogenesis.

Despite an increase in cell volume, previous results suggest that cells do not become polyploid during the 0-18h APF window^10^, raising the question of how cellular growth is achieved. There is growing evidence that ploidy sets an upper limit on physiological cell growth^38^. Recent findings in cultured cells indicate that cell volume increase is higher in G2 than in G1^39^. In this context, a G2 arrest at the L/P transition could facilitate cell growth in the absence of proliferation. Interestingly, we previously showed that random variations in tissue size are buffered during a specific time-window at the beginning of the pupal stage^20^. Our current findings imply that size adjustment occurs while the tissue is still actively growing, emphasizing the importance of this newly identified growth phase for final organ size determination.

Using inducible genetic systems, we triggered perturbations of growth-regulatory pathways in a careful time-controlled manner during early pupal development. Despite strong effects observed in adult wings, we found that the morphogens Wg and Dpp are dispensable for pupal wing growth. This suggests that morphogens are involved in non-growth-related processes during pupa that influence adult wing area. Of note, Dpp is required during pupal development to sustain the size-reductive cell divisions occurring between 16-24h APF^25^, although the biological significance of these divisions remains unclear. Reducing cell size at this stage could be crucial to ensure proper cellular dynamics underlying subsequent tissue morphogenesis such as hinge contraction, wing folding and unfolding^28,40^.

Our data highlight the importance of two nutrient-responsive pathways, IIS and TORC1, in regulating growth after the L/P transition. Further investigation is needed to understand how these pathways are activated during the non-feeding pupal stage to sustain imaginal disc development. A recent study established that the temporal allocation of amino acid resources between larval and pupal periods drives growth of pupal tissues^41^. In this view, the reuptake and processing of storage proteins by fat cells at the L/P transition, mediated by the hexamerin-like Fbp1, could provide a source of amino acids and trigger TORC1 activation in pupal tissues during the non-feeding phase. Consistent with this, levels of free amino acids rise in early pupa and remain elevated throughout metamorphosis^42^.

IIS also plays a conserved role in adjusting growth and metabolism in response to nutrient supply^43^. In *Drosophila*, IIS is activated by extracellular ligands that belong to the family of Dilps (*Drosophila* insulin-like peptides). While most Dilps are controlled at the secretion level by dietary amino acids during the larval stages, *dilp6* is transcriptionally induced under non-feeding conditions by fat body cells and plays a role during the pupal stage for proper adult body size^44,45^. High levels of *dilp6* expression in early pupal development could therefore act as relay to maintain IIS levels. However, whether physiological modifications of IIS and TORC1 activities serve as a timer for growth arrest, and what the limiting factor(s) could be, remains to be established.

Finally, our results suggest that ecdysone signaling is not required for the regulation of pupal wing growth. This result contrasts with the loss-of-function phenotypes during larval development, pointing to a specific role for the hormone in sustaining proliferative growth at that stage ^30–33^. Interestingly, expression of *fbp1* and of *dilp6* by fat body cells at the L/P transition is triggered by the peak of ecdysone^41,44^. Therefore, ecdysone could exert an indirect growth-promoting role at and after the L/P transition through the induction of Dilp6 and Fbp1.

In conclusion, our analysis of tissue growth during wing development reveals a previously unknown phase of non-proliferative growth during the early pupal stage, with significant implications for size adjustment and early morphogenesis. Importantly, this study redefines the timing of growth arrest, a key point for future research aimed at investigating the mechanisms that regulate the cessation of growth.

## RESOURCE AVAILABILITY

### Lead contact

Requests for further information and resources should be directed to and will be fulfilled by the lead contact, Laura Boulan.

### Materials availability

All reagents generated in this study are available from the lead contact.

### Data and code availability

This study did not generate data sets or codes.

## ACKNOWLEDGEMENTS

We thank David Lubensky and all lab members for insightful discussions. We thank Yohanns Bellaïche, Florencia di Pietro, Floris Bosveld, Marco Milán and Michael O’Connor for reagents; Aya Alami, Bénédicte Leveillé-Nizerolle and Sunitha Narasimha for technical help; the Bloomington Stock Center and the Vienna Drosophila RNAi Center for fly stocks; the PICT-IBiSA@BDD light-microscopy facility of Institut Curie; the MRI and Droso platforms from BioCampus Montpellier. This work was supported by CRBM, Institut Curie, CNRS, INSERM, FRM, the ATIP-Avenir program (starting grant to L.B.), the European Research Council (Advanced Grant # 694677 to P.L.), the Labex DEEP program (ANR-11-LABX-0044, ANR-10-IDEX-0001-02), PSL research University (Ph.D. fellowship to K.E.M.), and the Human Frontier Science Program (RGP0031/2020 to P.L.).

## AUTHOR CONTRIBUTIONS

Conceptualization, L.B. and P.L.; Methodology, K.E.M., I.G., E.D.G. and L.B.; Investigation, K.E.M., I.G., E.D.G. and L.B.; Writing – Original Draft, L.B.; Writing – Review & Editing, L.B. and P.L.; Funding Acquisition, L.B. and P.L.; Supervision, L.B.

## SUPPLEMENTAL INFORMATION

Document S1. Figures S1-S3

## EXPERIMENTAL PROCEDURES

### *Drosophila* lines and maintenance

Animals were reared at 25 °C (unless otherwise stated in the figure legends or other Methods sections) on fly food containing, per liter: 7.5g agar, 35g wheat flour, 50g yeast powder, 55g sugar, 25ml Methyl, and 4ml propionic acid.

The following lines were used: *rn-GAL4* (BDSC 7405); *UAS-GFP* (BDSC 35786); *w*^*1118*^ (BDSC 3605); *UAS-EcR*^*DN*^ (F645A, BDSC 6869); *UAS-EcR*^*RNAi*^ pan (BDSC 29374); *UAS-2xGFP* (BDSC 6874); *UAS-PTEN* (BDSC 82170); *UAS-Rheb*^*RNAi*^ (BDSC 33966); *UAS-Rheb*^*AV4*^ (BDSC 9690); *UAS-EGFR*^*CA*^ (BDSC 9533); *UAS-InR*^*DN*^ (BDSC 8252); *UAS-InR*^*CA*^ (BDSC 8263); *UAS-Akt*^*RNAi*^ (BDSC 33615); *tub-GAL80*^*ts*^ (BDSC 7017); *UAS-PTEN*^*RNAi*^ (NIG-Fly 5671); *nab-GAL4* (DGRC Kyoto 104533); *UAS-TSC1*^*RNAi*^ (VDRC 110811); *UAS-EGFR*^*RNAi*^ (VDRC 107130); *nub-GAL4*^46^; *UAS-EcI*^47^; *UAS-TSC1,UAS-TSC2 (UAS-TSC1/2)*^48^; *UAS-Cyp18A1*^49^. *UAS-notum* and *UAS-brk* were kindly provided by Marco Milán; *hsflp;UAS-PH-GFP;act>cd2>GAL4* was kindly provided by the Bellaïche lab; *elav-GAL80* (2) was kindly provided by Alex Gould. *rn-ShineGAL4* was generated in this work (see below).

### Generation of the *rn-ShineGAL4*

The *rn-ShineGAL4* stock is a light-inducible split-GAL4 composed by two transgenes: rn-Gal4DBD:2xnMagHigh1 recombined with ubi-2xpMag:AD-p10 (kind gift from Yohanns Bellaïche and Florencia di Pietro^29^). The rn-Gal4DBD:2xnMagHigh1 line was generated by gene conversion of the *rn-GAL4* driver (as described in^29^). It is composed by two tandem repeats of negative Magnet High1 variant photoswitches (nMagHigh1) fused to the GAL4 DNA binding domain (Gal4DBD: 1– 147 aa) under the control of the *rotund* promoter. The classical *rn-GAL4* line was crossed with *nanos-Cas9* (yw;nos-Cas9(II-attP40)). The transheterozygous F1 embryos were injected with pTV Gal4DBD:2xnMagHigh1 and pCFD4-Gal4T to replace the GAL4 sequence with the sequence of Gal4DBD:2xnMagHigh1, and resulting individuals were selected using 3xP3-Cherry.

### Samples preparation and processing

Only fixed tissues obtained from females were analyzed in this study. Between 0 and 8h APF, pupae were opened with dissection scissors first on the posterior side to remove the pressure, then a dorsal cut was performed to open the animal from posterior to anterior (at that stage the wing precursors are still in a ventro-lateral position). After 16h APF, the pupal case was carefully removed at the anterior and posterior extremities to access animal tissues. A small cut was then performed on the posterior and anterior parts of the animals to allow entry of the fixative. After fixation, animals were placed back to PBS, the remaining pupal case was removed, and the wings were carefully detached from the body. Finally, the animal’s cuticle around the wings could be removed to access the tissue. Fixation with 4% formaldehyde (Sigma) was performed at room temperature for 30min for samples until 8h APF, and for 1h30 for later pupal stages. After immunostaining^20^, tissues were mounted without coverslips on cell culture dishes with glass bottom (“Cellview”; Greiner Bio-one, #627861), in order to preserve the 3D structure. A Zeiss LSM900 Inverted Laser Scanning Confocal Microscope was used to image the samples and Z-stacks were processed using the FIJI and Imaris softwares.

Primary antibodies are the following: rabbit anti-fibrillarin (Abcam ab5821; 1/500), mouse anti-Lamin Dm0 (DSHB ADL84.12; 1/50).

### Volume measurements and growth rate analysis

Surfaces of the GFP signal and tissue volume were obtained using Imaris as previously described^20^. The growth rate k(t) based on tissue volume (V) is given by 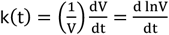. We approximated the derivatives for our data using 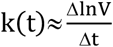.

### Quantification of cell number and volumes

At the tissue scale, nucleolar staining was used as a proxy to label nearly all cells with a single dot. The Spots detection function in Imaris allowed counting total cell number and deducing average cell volume from tissue volume (Figures 2A-B). For the clonal analysis (Figure 2C), *hsflp;UAS-PH-GFP;act>cd2>GAL4* animals were heat shocked for 5 minutes in mid-late larval development and synchronized at the L/P transition. Using Imaris, clones were outlined based on the PH-GFP membrane signal, and the inside of the cells was filled using the ‘Fill Holes’ function in Fiji before final volume measurement.

### Analysis of adult wings

Adult female flies of the appropriate genotypes were collected, stored in ethanol and mounted in a lactic acid:ethanol (6:5) solution. Pictures were acquired with a 1024 × 768 resolution using a MZ16-FA Leica Fluorescence Stereomicroscope with a DFC-490 Leica digital camera, and wing area was measured using automated segmentation as previously described^20^.

### Developmental timing measurement

Eggs were collected every 4h on plates made of 2% agar and 2% sucrose in PBS. The day after, L1-stage animals were synchronized and transferred to tubes with regular fly food. The number of new pupae was scored at the indicated times until all larvae pupariated.

### Statistics

For comparisons of the means between different genotypes, ANOVA tests were performed using GraphPad Prism 10.

### Shifts for GAL4 induction in early pupae

For temperature shifts, a tester line bearing *elav-GAL80, nub-GAL4, tub-GAL80*^*TS*^ was crossed to the UAS transgenes of interest. After short egg laying, larvae were kept at 18°C for 9 days. At the wandering stage, tubes were transferred to 29°C. After 5h, all pupae were removed to ensure that all animals have activated enough GAL4 at the L/P transition. 3 hours later, newly pupariated animals were isolated in a separate tube and kept at 29°C for 16 hours. Animals were then placed back at 18°C until adult hatching. A similar approach was used for the *ShineGAL4* system: embryos were placed in the dark (at 25°C) after egg laying until the wandering stage, when they were transferred to constant daylight. Animals were synchronized at the L/P transition like for the TS system, adding that females were selected among newly pupariated animals. After 20h under the light, pupal wings were dissected, fixed and analyzed.

---

**Supplementary Figure 1.**
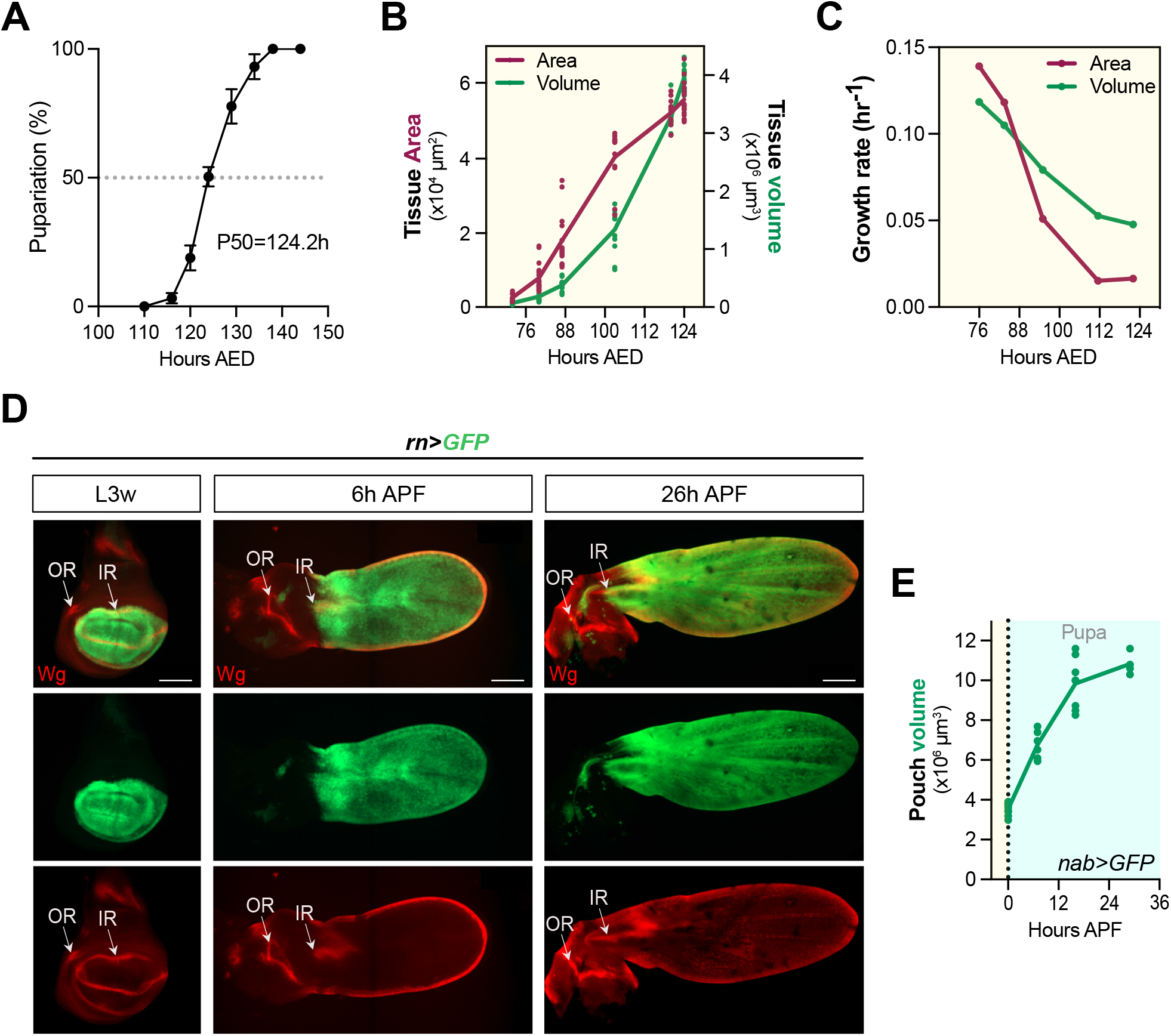
A) Pupariation curve of *rn>GFP* larvae at 25°C. B) Comparison of the growth curves during the last larval stage obtained by measuring either the area of the wing pouch (2D projection) or the tissue volume. The same samples were used for the two curves (and in Figure 1). C) Growth rate calculations based on the data presented in B. D) Comparison of the *rn>GFP* domain with Wingless (Wg) expression used as a landmark to show that the domain of GFP expression remains restricted to the future wing blade delimited by the IR of Wg. IR: Inner ring; OR: Outer ring. E) Growth curve in the early pupa based on GFP expression driven by another wing-specific driver: *nab-GAL4*. AED: After egg deposition; APF: After pupa formation.

**Supplementary Figure 2.**
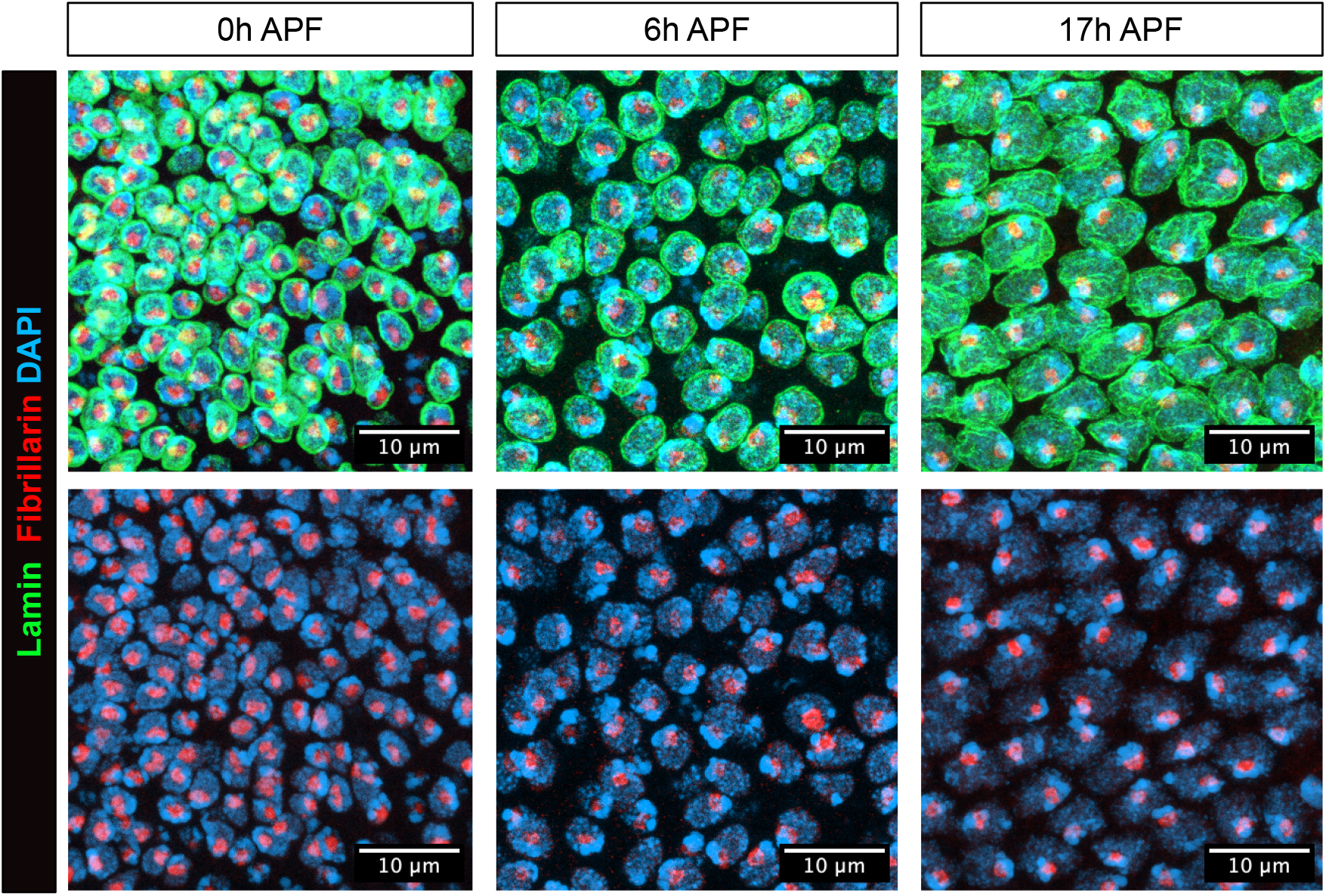
Nucleolar staining is used to label individual wing cells with a single dot, allowing to count cell number in a region of interest. Here are shown representative examples of wing primordia at the indicated time points highliting the nucleolus (Fibrillarin), the nuclear envelope (Lamin) and the DNA (DAPI).

**Supplementary Figure 3.**
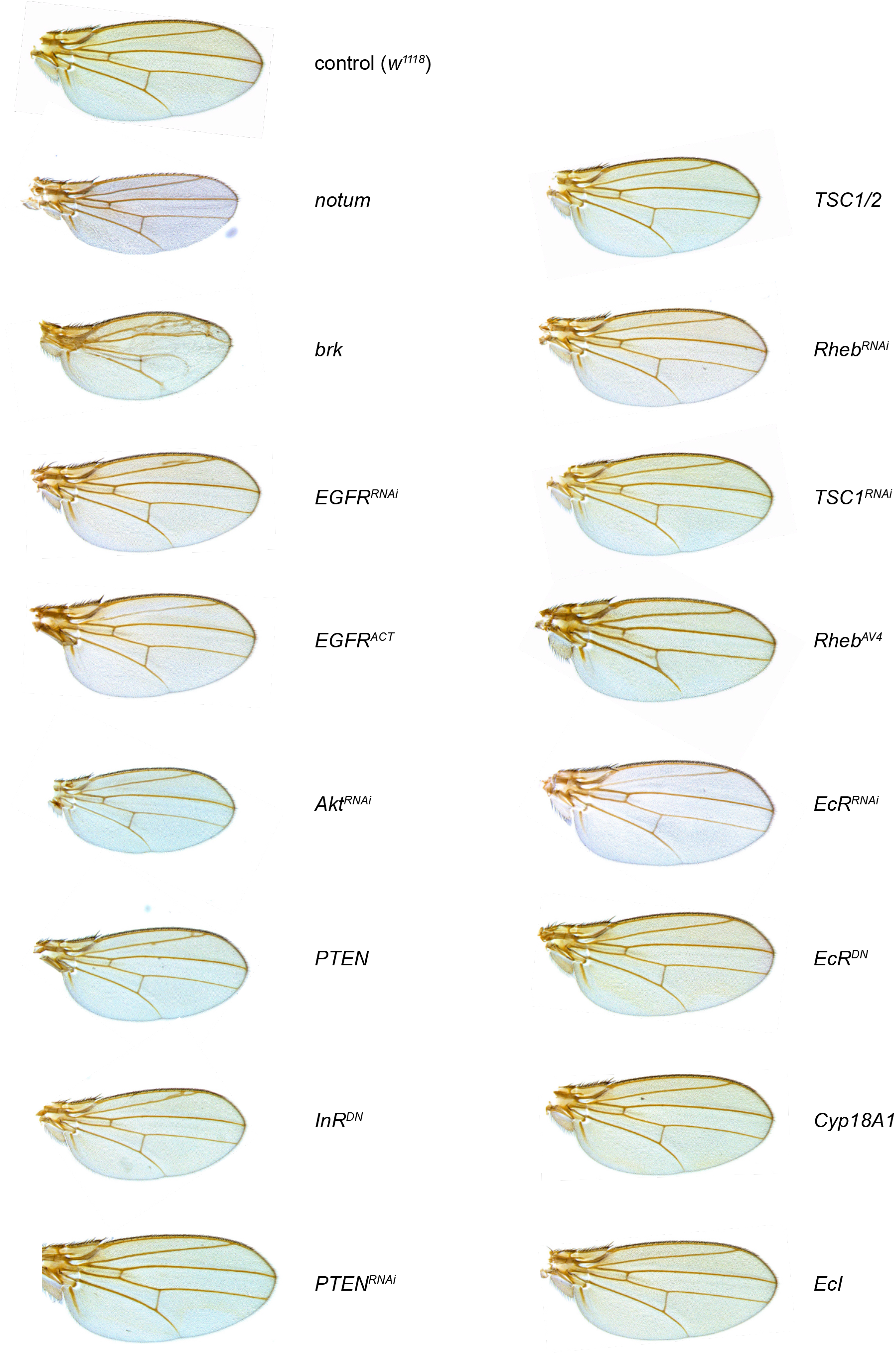
Representative examples of wings from animals carrying the *tub-GAL80TS, elavGAL80, nub-GAL4* driver combined with the indicated UAS transgenes. Activation of the GAL4/UAS system in the wing was specifically triggered during the 0-16h APF time window using a shift at 29°C.

## REFERENCES

1. Dekanty, A., and Milán, M. (2011). The interplay between morphogens and tissue growth. EMBO reports 12, 1003–1010. 10.1038/embor.2011.172.

2. Boulan, L., Milán, M., and Léopold, P. (2015). The Systemic Control of Growth. Cold Spring Harbor Perspectives in Biology 7, a019117. 10.1101/cshperspect.a019117.

3. Boulan, L., and Léopold, P. (2021). What determines organ size during development and regeneration? Development (Cambridge, England) 148. 10.1242/dev.196063.

4. Penzo-Méndez, A.I., and Stanger, B.Z. (2015). Organ-Size Regulation in Mammals. Cold Spring Harbor perspectives in biology 7, a019240. 10.1101/cshperspect.a019240.

5. Tripathi, B.K., and Irvine, K.D. (2022). The wing imaginal disc. Genetics 220, iyac020. 10.1093/genetics/iyac020.

6. Harmansa, S., and Lecuit, T. (2021). Forward and feedback control mechanisms of developmental tissue growth. Cells & Development 168, 203750. 10.1016/j.cdev.2021.203750.

7. Hariharan, I.K. (2015). Organ Size Control: Lessons from Drosophila. Developmental Cell 34, 255–265. 10.1016/j.devcel.2015.07.012.

8. Fain, M.J., and Stevens, B. (1982). Alterations in the cell cycle of Drosophila imaginal disc cells precede metamorphosis. Developmental Biology 92, 247–258. 10.1016/0012-1606(82)90169-5.

9. Milán, M., Campuzano, S., and García-Bellido, A. (1996). Cell cycling and patterned cell proliferation in the Drosophila wing during metamorphosis. Proceedings of the National Academy of Sciences 93, 11687–11692. 10.1073/pnas.93.21.11687.

10. Guo, Y., Flegel, K., Kumar, J., McKay, D.J., and Buttitta, L.A. (2016). Ecdysone signaling induces two phases of cell cycle exit in Drosophila cells. Biology Open 5. 10.1242/bio.017525.

11. Etournay, R., Popović, M., Merkel, M., Nandi, A., Blasse, C., Aigouy, B., Brandl, H., Myers, G., Salbreux, G., Jülicher, F., et al. (2015). Interplay of cell dynamics and epithelial tension during morphogenesis of the Drosophila pupal wing. eLife 4. 10.7554/eLife.07090.

12. Vollmer, J., and Iber, D. (2016). An Unbiased Analysis of Candidate Mechanisms for the Regulation of Drosophila Wing Disc Growth. Sci Rep 6, 39228. 10.1038/srep39228.

13. Wartlick, O., Mumcu, P., Kicheva, A., Bittig, T., Seum, C., Jülicher, F., and González-Gaitán, M. (2011). Dynamics of Dpp Signaling and Proliferation Control. Science 331, 1154–1159. 10.1126/science.1200037.

14. Nienhaus, U., Aegerter-Wilmsen, T., and Aegerter, C.M. (2012). In-Vivo Imaging of the Drosophila Wing Imaginal Disc over Time: Novel Insights on Growth and Boundary Formation. PLoS ONE 7.

15. Hufnagel, L., Teleman, A.A., Rouault, H., Cohen, S.M., and Shraiman, B.I. (2007). On the mechanism of wing size determination in fly development. Proceedings of the National Academy of Sciences of the United States of America 104. 10.1073/pnas.0607134104.

16. Aegerter-Wilmsen, T., Aegerter, C.M., Hafen, E., and Basler, K. (2007). Model for the regulation of size in the wing imaginal disc of Drosophila. Mechanisms of Development 124. 10.1016/j.mod.2006.12.005.

17. Aegerter-Wilmsen, T., Heimlicher, M.B., Smith, A.C., de Reuille, P.B., Smith, R.S., Aegerter, C.M., and Basler, K. (2012). Integrating force-sensing and signaling pathways in a model for the regulation of wing imaginal disc size. Development (Cambridge) 139.

18. Strassburger, K., Lutz, M., Müller, S., and Teleman, A.A. (2021). Ecdysone regulates Drosophila wing disc size via a TORC1 dependent mechanism. Nat Commun 12, 6684. 10.1038/s41467-021-26780-0.

19. Zhao, Y., Alexandre, C., Kelly, G., Perez-Mockus, G., and Vincent, J.-P. (2024). Growth-induced physiological hypoxia correlates with growth deceleration during normal development. Preprint at bioRxiv, https://doi.org/10.1101/2024.06.04.597345 10.1101/2024.06.04.597345.

20. Blanco-Obregon, D., El Marzkioui, K., Brutscher, F., Kapoor, V., Valzania, L., Andersen, D.S., Colombani, J., Narasimha, S., McCusker, D., Léopold, P., et al. (2022). A Dilp8-dependent time window ensures tissue size adjustment in Drosophila. Nat Commun 13, 5629. 10.1038/s41467-022-33387-6.

21. Harmansa, S., Erlich, A., Eloy, C., Zurlo, G., and Lecuit, T. (2023). Growth anisotropy of the extracellular matrix shapes a developing organ. Nat Commun 14, 1220. 10.1038/s41467-023-36739-y.

22. Kumar, N., Rangel Ambriz, J., Tsai, K., Mim, M.S., Flores-Flores, M., Chen, W., Zartman, J.J., and Alber, M. (2024). Balancing competing emects of tissue growth and cytoskeletal regulation during Drosophila wing disc development. Nat Commun 15, 2477. 10.1038/s41467-024-46698-7.

23. Barrio, L., and Milán, M. (2017). Boundary Dpp promotes growth of medial and lateral regions of the Drosophila wing. eLife 6. 10.7554/eLife.22013.

24. Barrio, L., and Milán, M. (2020). Regulation of Anisotropic Tissue Growth by Two Orthogonal Signaling Centers. Developmental Cell 52. 10.1016/j.devcel.2020.01.017.

25. Gui, J., Huang, Y., Montanari, M., Toddie-Moore, D., Kikushima, K., Nix, S., Ishimoto, Y., and Shimmi, O. (2019). Coupling between dynamic 3D tissue architecture and BMP morphogen signaling during Drosophila wing morphogenesis. Proceedings of the National Academy of Sciences 116, 4352–4361. 10.1073/pnas.1815427116.

26. Guittard, E., Blais, C., Maria, A., Parvy, J.-P., Pasricha, S., Lumb, C., Lafont, R., Daborn, P.J., and Dauphin-Villemant, C. (2011). CYP18A1, a key enzyme of Drosophila steroid hormone inactivation, is essential for metamorphosis. Developmental biology 349, 35–45. 10.1016/j.ydbio.2010.09.023.

27. Okamoto, N., Viswanatha, R., Bittar, R., Li, Z., Haga-Yamanaka, S., Perrimon, N., and Yamanaka, N. (2018). A Membrane Transporter Is Required for Steroid Hormone Uptake in Drosophila. Developmental Cell 47, 294-305.e7. 10.1016/j.devcel.2018.09.012.

28. Diaz de la Loza, M.C., and Thompson, B.J. (2017). Forces shaping the Drosophila wing. Mechanisms of Development 144, 23–32. 10.1016/j.mod.2016.10.003.

29. di Pietro, F., Herszterg, S., Huang, A., Bosveld, F., Alexandre, C., Sancéré, L., Pelletier, S., Joudat, A., Kapoor, V., Vincent, J.-P., et al. (2021). Rapid and robust optogenetic control of gene expression in Drosophila. Developmental Cell 52.

30. Delanoue, R., Slaidina, M., and Leopold, P. (2010). The steroid hormone ecdysone controls systemic growth by repressing dMyc function in Drosophila fat cells. Dev Cell 18, 1012–1021. https://doi.org/S1534-5807(10)00215-7 [pii] 10.1016/j.devcel.2010.05.007.

31. Herboso, L., Oliveira, M.M., Talamillo, A., Pérez, C., González, M., Martín, D., Sutherland, J.D., Shingleton, A.W., Mirth, C.K., and Barrio, R. (2015). Ecdysone promotes growth of imaginal discs through the regulation of Thor in D. melanogaster. Scientific Reports 5, 12383. 10.1038/srep12383.

32. Dye, N.A., Popović, M., Spannl, S., Etournay, R., Kainmüller, D., Ghosh, S., Myers, E.W., Jülicher, F., and Eaton, S. (2017). Cell dynamics underlying oriented growth of the Drosophila wing imaginal disc. Development (Cambridge, England) 144, 4406–4421. 10.1242/dev.155069.

33. Perez-Mockus, G., Cocconi, L., Alexandre, C., Aerne, B., Salbreux, G., and Vincent, J.-P. (2023). The Drosophila ecdysone receptor promotes or suppresses proliferation according to ligand level. Developmental Cell 58, 2128-2139.e4. 10.1016/j.devcel.2023.08.032.

34. O’Farrell, P.H. (2004). How Metazoans Reach Their Full Size: The Natural History of Bigness. Cold Spring Harbor Monograph Archive 42, 1–22. 10.1101/0.1-22.

35. Cadart, C., Bartz, J., Oaks, G., Liu, M.Z., and Heald, R. (2023). Polyploidy in Xenopus lowers metabolic rate by decreasing total cell surface area. Current Biology 33, 1744-1752.e7. 10.1016/j.cub.2023.03.071.

36. Neufeld, T.P., de la Cruz, A.F., Johnston, L.A., and Edgar, B.A. (1998). Coordination of growth and cell division in the Drosophila wing. Cell 93, 1183–1193.

37. Weigmann, K., Cohen, S.M., and Lehner, C. (1997). Cell cycle progression, growth and patterning in imaginal discs despite inhibition of cell division after inactivation of Drosophila Cdc2 kinase. Development 124, 3555–3563.

38. Cadart, C., and Heald, R. (2022). Scaling of biosynthesis and metabolism with cell size. MBoC 33, pe5. 10.1091/mbc.E21-12-0627.

39. Cadart, C., Venkova, L., Piel, M., and Cosentino Lagomarsino, M. (2022). Volume growth in animal cells is cell cycle dependent and shows additive fluctuations. eLife 11, e70816. 10.7554/eLife.70816.

40. Hadjaje, S., Andrade-Silva, I., Dalbe, M.-J., Clément, R., and Marthelot, J. (2024). Wings expansion in Drosophila melanogaster. Preprint, https://doi.org/10.1101/2024.05.17.594352 10.1101/2024.05.17.594352.

41. Valzania, L., Alami, A., and Léopold, P. (2024). A temporal allocation of amino acid resources ensures fitness and body allometry in Drosophila. Developmental Cell. 10.1016/j.devcel.2024.05.018.

42. An, P., Yamaguchi, M., and Fukusaki, E. (2017). Metabolic profiling of Drosophila melanogaster metamorphosis: a new insight into the central metabolic pathways. Metabolomics 13, 1–13. 10.1007/s11306-017-1167-1.

43. Suzawa, M., and Bland, M.L. (2023). Insulin signaling in development. Development 150, dev201599. 10.1242/dev.201599.

44. Slaidina, M., Delanoue, R., Gronke, S., Partridge, L., and Léopold, P. (2009). A Drosophila insulin-like peptide promotes growth during nonfeeding states. Developmental cell 17, 874–884. 10.1016/j.devcel.2009.10.009.

45. Okamoto, N., Yamanaka, N., Yagi, Y., Nishida, Y., Kataoka, H., O’Connor, M.B., and Mizoguchi, A. (2009). A fat body-derived IGF-like peptide regulates postfeeding growth in Drosophila. Developmental cell 17, 885–891. 10.1016/j.devcel.2009.10.008.

46. Azpiazu, N., and Morata, G. (2000). Function and regulation of homothorax in the wing imaginal disc of Drosophila. Development 127, 2685–93.

47. Okamoto, N., Viswanatha, R., Bittar, R., Li, Z., Haga-Yamanaka, S., Perrimon, N., and Yamanaka, N. (2018). A Membrane Transporter Is Required for Steroid Hormone Uptake in Drosophila. Developmental Cell 47, 294-305.e7. 10.1016/j.devcel.2018.09.012.

48. Tapon, N., Ito, N., Dickson, B.J., Treisman, J.E., and Hariharan, I.K. (2001). The Drosophila tuberous sclerosis complex gene homologs restrict cell growth and cell proliferation. Cell 105, 345–355.

49. Rewitz, K.F., Yamanaka, N., and O’Connor, M.B. (2010). Steroid hormone inactivation is required during the juvenile-adult transition in Drosophila. Developmental cell 19, 895–902. 10.1016/j.devcel.2010.10.021.

